# Mysterious disappearances of a large mammal in Neotropical forests

**DOI:** 10.1101/2020.12.08.416552

**Authors:** J.M.V. Fragoso, A.P. Antunes, M. Altrichter, P.A.L. Constantino, G. Zapata-Ríos, M. Camino, B. de Thoisy, R.B. Wallace, H.R. El Bizri, T.Q. Morcatty, P. Mayor, C. Richard-Hansen, M.T. Hallett, R.A. Reyna-Hurtado, H. Beck, S. de Bustos, R.E. Bodmer, A. Keuroghlian, A. Nava, O.L. Montenegro, E. Painkow Neto, A.L.J. Desbiez, K.M. Silvius

## Abstract

The drivers of periodic population cycling by some animal species in northern systems remain unresolved^1^. Mysterious disappearances of populations of the Neotropical, herd-forming white-lipped peccary (*Tayassu pecari*, henceforth “WLP”) have been anecdotally documented and explained as local events resulting from migratory movements or overhunting^2,3,4^, or as disease outbreaks^5,6^, and have not been considered in the context of large-scale species-specific population dynamics. Here we present evidence that WLP disappearances represent troughs in population cycles that occur with regular periodicity and are synchronized at regional and perhaps continent-wide spatial scales. Analysis of 43 disappearance events and 88 years of commercial and subsistence harvesting data reveals boom – bust population cycles lasting from 20 to 30 years, in which a rapid population crash occurring over 1 to 5 years is followed by a period of absence of 7 to12 years and then a slow growth phase. Overhunting alone cannot explain the crashes, but as in northern systems dispersal during the growth phase appears to play a key role. This is the first documentation of population cycling in a tropical vertebrate.

## Main text

The white-lipped peccary (*Tayassu pecari*, henceforth “WLP”), is the only tropical forest mammal to form large, permanent, cohesive herds comprising hundreds of individuals. While the WLP’s unusual social organization is well-known, another characteristic of this keystone species—the disappearance and eventual reappearance of entire populations over large spatial scales—is only anecdotally documented and remains poorly understood.

The WLP has tremendous impact on forest composition and on the livelihoods of forest-dependent peoples. In the Amazon, the home range of a typical herd of about 400 animals may extend to 200 km^2(6)^. Considering an adult weight of 30 to 50 kg, such a herd represents 12,000 to 20,000 kg of biomass moving in synchrony, rooting up soils, consuming seeds, seedlings, plant parts and animal matter, dropping excreta, creating wallows, and serving as preferred prey for large cats, all factors that influence biotic communities^7^ and the carbon cycle^8^. WLP population dynamics are therefore of interest in studies of forest ecology, as periodicity in their abundance may permit periodic recruitment of plants and shifting predator pressure on other prey species. The WLP is also highly prized as food by indigenous peoples^9^ and plays a prominent role in their religious beliefs^10,11,12^. WLPs can contribute ≥ 70% of all animal biomass consumed in indigenous Yanomami and Kaxinawa villages^6,13^. Loss of WLP populations alters religious practices^12^ and may induce human emigration^14^.

In the 1980s, researchers in the Amazon began to report occasional and local disappearances of WLP populations, attributing them to large-scale movements^2,3^, overhunting^4^, or disease outbreaks^15^. In 1997, Fragoso^5^ documented the absence and subsequent reappearance of WLPs over a region the size of Norway, and suggested that epidemics could cause disappearances over very large, regional scales (millions of ha). No data were available to inform whether these events were related to local, novel disease outbreaks, as documented for temperate and tropical Old-World ungulates^16,17^, or an intrinsic and recurrent consequence of the species’ unique social organization, herd size and movements. Because WLPs are notoriously difficult to study in their remote, dense forest habitats, we fill this gap in knowledge by compiling multiple sources of evidence on fluctuations in WLP abundance throughout the species’ range and analyzing the extent, timing and periodicity of documented disappearances.

### Literature review and expert reporting of disappearances

We conducted a questionnaire-based survey of experts through 2019 and reviewed the literature on WLP disappearances. We recorded 67 reports from experts on 43 independent WLP disappearances at 38 sites in 9 countries (S1, S2), occurring over a cumulative area of at least 50 million ha. Twenty-eight disappearance sites are in contiguous old growth forest in the Amazon region, nine in forest fragments in Atlantic Forest in southern Brazil and northern Argentina, and one in rainforest in Guatemala (S1, S2).

At small scales (<100,000 ha), we defined a disappearance as a sudden (from one 12-month period to the next) decline from previously documented (henceforth “normal”) abundances to low abundance or absence. Over large areas and in the absence of intensive searching it is difficult to distinguish between low abundance and absence. We defined a disappearance as occurring when no or very few WLP are sighted for at least 3 years, and a reappearance as a return to normal population abundance for that region. For example, a density of 5 to 10 individuals per km^2^ is low to normal in the Amazon^18,19^, where high densities reach 100 or more individuals per km^2 (20)^. Not all population declines had been documented continuously; in these cases, the absence periods were estimated by researchers. Regional or large scale (>1,000,000 ha) disappearances may occur gradually over 2 to 5 years, with the start of the event progressing outward from one area^21^.

Two cases, one in Brazil and one in Guyana, illustrate rapid population crashes (S1). At Maracá Island Ecological Reserve in the northern Brazilian Amazon state of Roraima, WLPs were at high density (138.8 ind/km^2^) in 1988 and herds were sighted almost every day during intensive searching^20^. WLPs then disappeared in March 1989 and were not detected despite continuous monitoring until two herds were seen in 1991^6^. Herd size grew very slowly and by 2005 the population reached normal to high density. In Guyana, intensive research over 3.5 years described a WLP population with high densities up until June 2010, when this study ended^9,14^. WLPs then disappeared over the course of one year and their absence was documented starting in 2011 by camera trapping, transect surveys and hunter reports for the following 7 years (S2). They had not reappeared as of 2019 in most areas.

Population absences or very low abundances for 23 cases in contiguous forest lasted on average 10.3 years (SD = 4.9), with 50% lasting between 7 and 12 years. Some populations lows were well documented through long-term field research. For example, in contiguous forest regions of the Amazon biome in French Guiana, WLPs were continuously monitored between 2000 and 2019. They were at normal abundances throughout the state in 2000 and 2001. Monitoring revealed a geographically expanding disappearance starting in 2003, reaching almost complete absence from 2006 to 2009 across the entire region. During 2012 and 2013, WLPs reappeared in several locations.

Longer lasting disappearances are reported from discontinuous forests outside the Amazon. In the 41,704-ha Intervales State Park, an Atlantic Forest fragment in São Paulo State, WLPs disappeared in 1990 and reappeared in 2017 after a 27-year absence. In the larger Iguaçu National Park and adjacent northern Misiones province in Argentina, WLPs disappeared around 1995. Years of surveys confirmed their absence until several individuals were detected with camera traps in 2016, likely immigrating from nearby forested areas in Argentina^22^. In Turvo State Park (Rio Grande do Sul State, Brazil), WLPs disappeared in 1980 and have not been seen since then (S1).

More than one disappearance event has been documented through time at several sites, pointing to periodicity in the phenomenon. In and around Manu National Park, Peru, WLPs disappeared in 1978, reappeared 12 years later, then disappeared again in 2011. A few individuals were detected in 2015, but as of 2019, the population had not returned to pre-disappearance levels. Two distinct disappearance events occurred in the 275,531-ha Jutaí River Extractive Reserve, located in continuous, undisturbed forest in western Amazonas state, Brazil. Local inhabitants reported the start of the first disappearance between 1998-2002, with the timing depending on the location within the reserve. WLPs reappeared between 2006-2011, with the timing again depending on the site (S2). Researchers were present for the second disappearance, which began in 2013 near the mouth of the river (S2). By 2014 WLPs had also disappeared from the adjacent section of the watershed and in 2015 from the more distant headwater region. WLPs have not yet been detected at any location in the reserve. In Madidi National Park, Bolivia, WLPs disappeared in 1984 for 10 to 14 years, then reached normal densities for several years, and disappeared again in mid-2017 along the Heath river, followed by the Tuichi and Hondo rivers in 2018 (S2). Three disappearance events were reported in an area of over 3 million ha, covering the Amanã Sustainable Development Reserve, Unini Extractive Reserve and Jaú National Park (S2).

Because researchers monitor a limited set of sites in a region, the true geographical extent of disappearances is usually not known. Nevertheless, we documented several cases of disappearances occurring at massive scales: 35% of the disappearances took place over areas larger than 1-million ha (S1). In French Guiana, WLPs disappeared over the entire 8-million ha territory (Fig. 1). In Guyana, they disappeared over at least 5.4-million ha in the Rupununi region. In the northern Amazon of Brazil, WLPs disappeared over at least 4.8-million ha in Roraima state (Fig. 1). Madidi National Park in Bolivia and Manu National Park in Peru experienced disappearances over at least about 1-million and 1.7-million ha, respectively (S2).

**Fig. 1.**
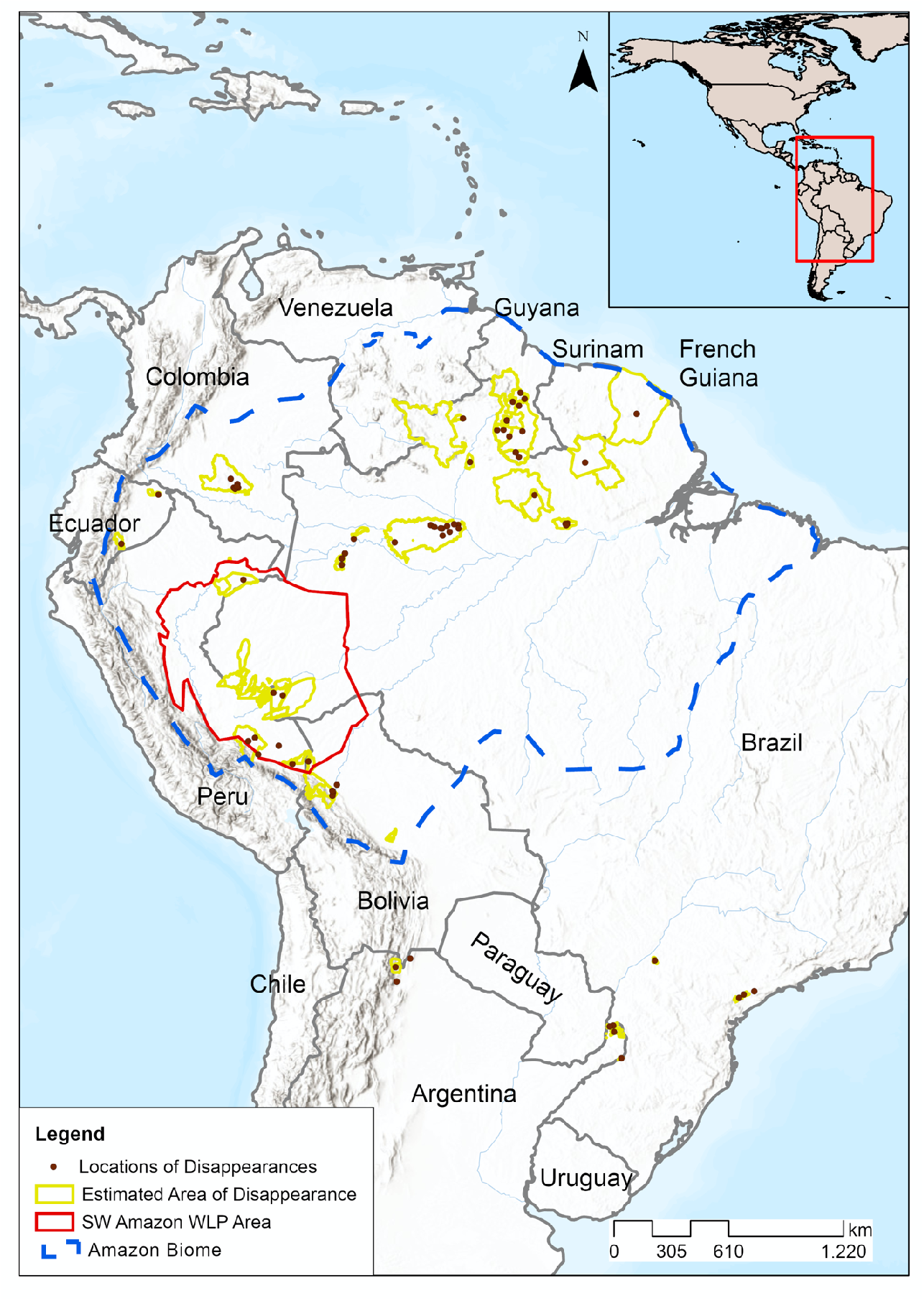
Locations of white-lipped peccary (Tayassu pecari) disappearances. Brown dots mark disappearances of small or unknown areal extent. Yellow lines mark the estimated extent for disappearances over larger regions. The 686 million ha SW Amazon region, the source area for pelt and hunting data, is delineated in red, and the blue line demarcates the Amazon biome. The disappearance site in Guatemala is not shown.

### Subsistence hunting and commercial pelt returns

We examined legal pelt trade records from 1932 to 2019^23,24^ and subsistence hunting from 66 studies between 1965 and 2017 (S3) from the Southwestern (SW) Amazon. The combined dataset includes records of almost 2 million WLPs hunted for their hides and meat over an estimated 686 million ha over 88 years (Fig. 1). We accounted for variation in hunting effort by calculating the ratio of WLP to collared peccary (*Pecari tajacu*) pelts and kills, then used the ratio of WLPs hunted to the two-species total to detect fluctuations in WLP peccary returns independent of overall hunting effort. The two peccary species are among the most preferred species by subsistence hunters^9,25^ and prices for their skins were nearly identical during the analysed period^23^. They differ in their socio-ecology; collared peccaries are territorial, less gregarious and form only small family groups (6 to 20 individuals)^26^. We modeled the fluctuations in the WLP ratio as a population fluctuation (see Online Methods section).

Declines in WLP abundance occurred synchronously over the SW Amazon (Fig. 1). At least three large abundance variations occurred through time (Fig. 2). Two peaks (1941 and 2006) were followed by rapid declines (late 1940s and late 2000s), while a third peak (1975) showed a more gradual decline, perhaps due to limited data during this period. Increases were gradual—population growth required over 20 years to reach population maxima—whereas the steeper slopes of two declines demonstrate large-scale, fast population crashes reaching minima after about 5 years over this huge area.

**Fig. 2.**
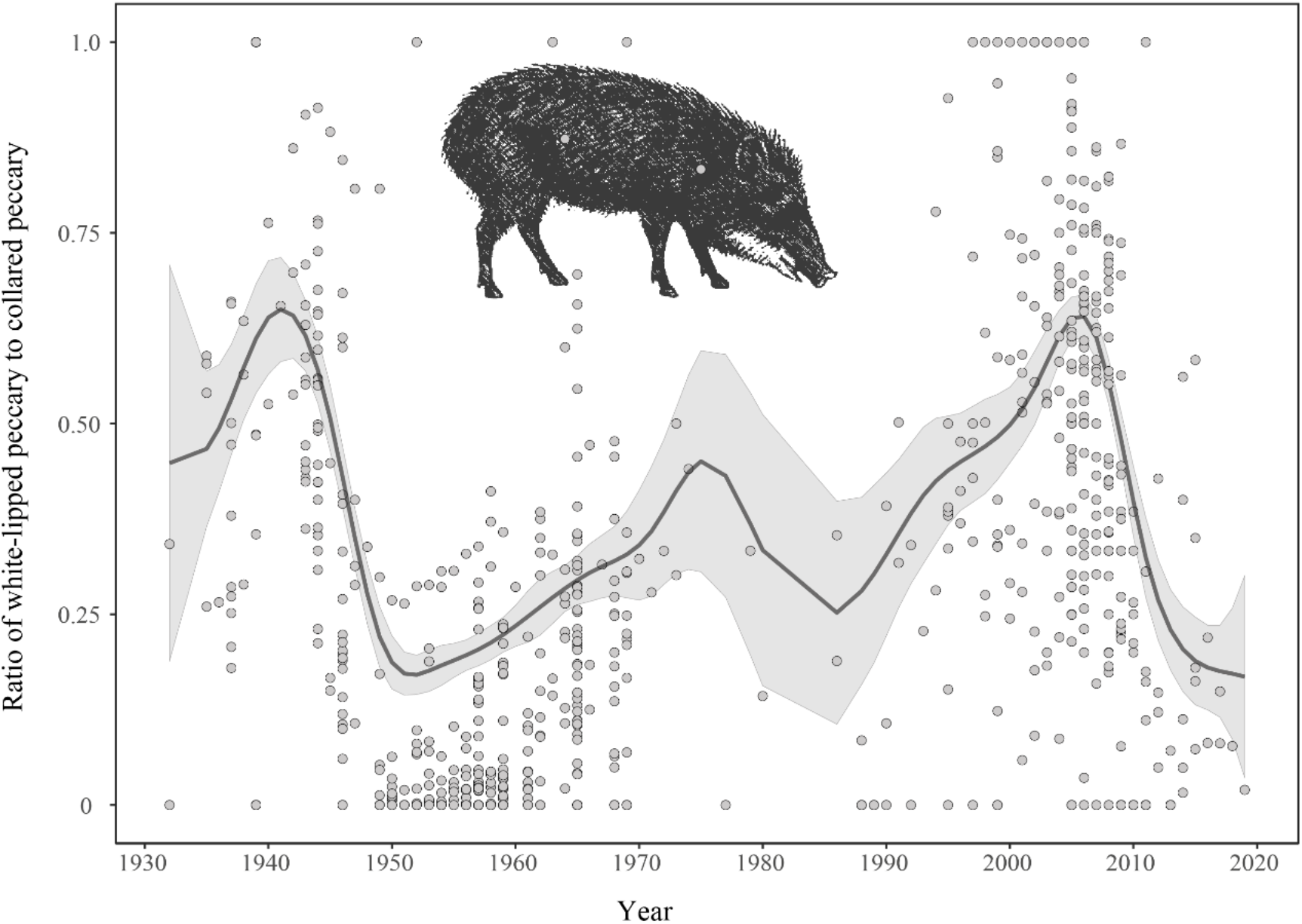
Estimated multi-decadal population cycles of WLP based on the ratio of hunted WLP to hunted collared peccary (1932 to 2019) in the southwestern Amazon. The shaded gray area indicates 95% confidence interval. Circles are individual data points. Data from commercial pelt and subsistence hunting for meat.

### WLP population cycles

The compiled data from multiple sources on timing and spatial extent of WLP disappearance events and abundances reveal cycling in WLP populations at local, landscape and perhaps even continental scales, documenting an enigmatic population dynamic pattern for one of the most iconic Neotropical species, and one that is unexpected for any large forest mammal. WLP population cycles appear to last from 20 to 30 years from peak to peak, with a rapid population decline over 1 to 5 years, and an absence or low-abundance phase of 7 to 12 years followed by slow growth over about 20 years. Such population cycling is previously undocumented for any tropical mammal. Ungulate population dynamics in the temperate zone have been shown to be influenced by weather, interspecific interactions, and food resources^27,28,29^, but none of these exhibit cycles or even extreme highs and lows over time as documented here for WLPs. Although the WLP is highly susceptible to overhunting^30^, these population cycles are unlikely caused by hunting—most disappearances occurred in unpopulated, remote areas of contiguous forest.

Although our data are restricted to recent history, WLP population cycling appears to have occurred for long enough through time to be incorporated into the cosmologies and beliefs of the indigenous people of the Amazon. Several ethnic groups describe disappearance events as linked to the death of a powerful shaman^6,12^. The shaman, in anger with his people, upon his death takes the WLPs into an underground cavern and blocks the entrance. Their return can only be secured by another shaman using special cultural activities to unblock the entrance. This suggests that population fluctuations are a characteristic of the species.

Periodicity in northern animal population dynamics has long been observed in nature^31^. However, the cause of population cycles has been a major unresolved issue in animal ecology for about a hundred years^1,31^. In general, cycling populations of vertebrates and insects show rapid decline phases and slow growth phases^1,32^, a pattern we document in our results. Research also indicates that three types of condition are necessary for a population to cycle^1,32^; two of these—high reproductive rates and the potential for overcompensating density dependent mortality are easily identifiable in WLPs, while the third—a negative condition prolonged enough to delay recovery—is not readily apparent and could be expected to vary across the species vast range from South to Central America. To date, reports of disappearances unrelated to overhunting come only from Amazon and Atlantic forest in South America, and not from the Pantanal or Cerrado or most Central American locations.

WLP’s have a relatively high reproductive rate (1.67 young per litter, 250-day intervals between litters, 18 months age of first reproduction)^33^. Their gregarious nature and unusually large herd size with overlapping herd home ranges may set the conditions for rapid disease spread that locally eliminates entire populations^34^. In an analogous situation, humans have had to decrease social interactions and maintain social-distancing to isolate and slow the transmission of the COVID-19 virus^35^.

WLP re-appearance consisting of gradually increasing numbers of individuals may result from population growth from a few remaining individuals or re-colonization^6^. Their obligate herding social structure may mean that populations cannot reach high growth rates until herds reach a threshold size; this, together with the need for immigration from other populations to re-establish local populations may be the factors that delay recovery, with variations in cycle length across the range influenced by external factors such as food availability or predator pressure. The period of absence or low abundance for WLP is around 7-12 years. The disappearance period was over 20 years in sites with high forest fragmentation due to anthropogenic impacts, suggesting that in the absence of dispersal, a population can become extinct. This supports the importance of dispersal in shaping the temporal and spatial scale of a cycle^1^.

The gregarious nature of WLP individuals within and amongst herds^26^ may allow higher amplification of pathogen transmission^34^. Populations of the sympatric collared peccary, which has small herds of 6-20 individuals and is territorial^26^, do not cycle or even fluctuate in a non-cyclical manner between extreme highs and lows. When disease plays a role in population cycling, the characteristics of the disease agent and vector also can influence the amplitude and periodicity of the cycle^1^. Emmons^15^ proposed a local disease outbreak for a WLP disappearance that occurred in Manu National Park, Peru, and Fragoso^5,6^ postulated epidemic outbreaks at regional scales after documenting a disappearance in northern Roraima, Brazil. Our study documented rapid declines in abundance, characteristic of catastrophic events such as disease in which individuals experience severe physiological stress that affect both maintenance and reproduction^1,36,37^. It is still an open question for WLPs whether disease outbreaks could drive populations cycles, on their own or in synchrony with other stressors.

WLP are notoriously difficult to study--their long-distance daily movements, and their aggressive behavior towards hunters and predators make them difficult to capture even under normal circumstances. Future efforts among a network of researchers and community collaborators, with improved technologies and the expansion of biodiversity monitoring programs in the most remote Amazonia regions, should be feasible to establish to finally understand the mysterious WLP disappearances and confirm the details and causes of the population cycles documented in this study.

## Online Methods

### Survey

In January 2016 we sent out a questionnaire to 58 neotropical wildlife researchers, including all members of the IUCN/SSC Peccary Specialist Group. To complement the questionnaire, we conducted a Google Scholar search of the key words white-lipped peccary, *Tayassu pecari* and peccary and extracted the same information as from survey respondents for instances of multiyear, extensive area disappearances. Independent reports from the same date and location were consolidated as one event, while separate reports from the same or adjacent locations within a year of each other were retained in the data set but counted as single events. Disappearances at sites separated by 100 km were considered distinct events. We estimated the mean years of absence, SD and IQR for cases at sites >100,000 ha that were surrounded by contiguous forest. In cases where a range of time was provided, we used the average of the range.

### Commercial pelt trade and subsistence hunting record

We examined legal pelt trade records from 1932 to 1969 from 379 cargo manifests, port registries and financial documents^23^, governmental statistics from Peru between 1946 and 2019^24^, and subsistence hunting records from 66 published and unpublished studies from 1965 to 2017 (S2) from the Southwestern Amazon. The combined dataset includes records of nearly 2 million WLPs hunted for their hides and meat over an estimated 686 million ha. We controlled for variation in hunting effort by calculating the ratio of WLP pelts and kills to other similarly preferred and priced species, the collared peccary (*Tayassu tajacu*). For each year and locality, we divided the number of WLP hunted by the total hunted of the two species. The resulted dataset contains 749 WLP hunting-proportions events for both commercial and subsistence purposes. We assume that the number of kills reflects the abundance of the species on the hunting grounds, and that when the number of hunted WLPs is low relative to collared peccary, the WLP populations are low and individuals are difficult to find and kill. Conversely, if the ratio of WLP to collared peccary is high, we assume that the WLP population is also high. Using the ratio of hunted WLP allowed the joint analysis of commercial and subsistence hunting data throughout time.

### Population fluctuations analysis

We modeled the fluctuations of the WLP ratio through time as population fluctuations. The unknown proportion of the WLPs hunted in relation to collared peccary in year *t*, for *t* = 1932 to 2019, denoted by *γ_i_*(*t*), is assumed to be a smooth curve over time. Since the proportion varied between 0 and 1 we used a beta inflated distribution model, *X*(*t*)~*BEINF*(*BI*(*t*), *α, γ, μ, ϕ*) – a mixture distribution between beta and Bernoulli distributions that allows values 0 and 1 for the response variable – where *BEINF*(*BI*(*t*)) denotes a beta inflated distribution with mean *BEINF*(*BI*(*t*)), *μ* location and *ϕ* scale parameters that control the probability density function, and two extra parameters *α* and *γ*, which model the probabilities at zero and one^38,39^. To track the population fluctuation curve *γ_i_*(*t*) as a cubic spline^40,41^, we used gamlss package (Version 5.1-6)^42^ in R language software^43^, which estimates *α, γ, μ, ϕ* and spline parameters (number and position of knots for the curve *γ_i_*(*t*)) by maximum likelihood with automatized Generalised Akaike Information Criteria for model selection^39^. We used the different Amazon river basins as a random factor due to different sample sizes among basins, and to possible non-independence of sampled localities within each basin.

## Acknowledgments

J.M.V.F. thanks the Yanomami people for documenting the return of WLP to their area in the 1990s, and the Makushi and Wapichan people for tracking their hunting returns so precisely. Thanks also to Roan McNab, Luis Pacheco, Mineração Rio do Norte and the Biota Projetos e Consultoria Ambiental for providing information. C.R.H. thanks the Parc Amazonien de Guyane and European Community funds for part of the French Guiana dataset. Thanks to Bill Magnusson and Albertina Lima for providing a home at INPA for the writing of a first draft of the manuscript.

## Author Contributions

J.M.V.F., R.R. and C.R.H. conceived the project. J.M.V.F wrote the first version of the manuscript. J.M.V.F., A.A, and P.A.L.C. designed the study. J.M.V.F., A.A, M.A., K.M.S., P.A.L.C., H.R.E.B. and G.Z-R. participated in analysis, and interpretation of data. All authors provided data. Subsequent versions of the manuscript were reviewed by all authors, with J.M.V.F., M.A. and K.M.S. incorporating suggested edits and rewriting as needed.

## Notes

### Competing Interest Statement

The authors have declared no competing interest.

## References

1. Myers, J. H. Population cycles: generalities, exceptions and remaining mysteries. Proc. R. Soc. Lond. B. 285, 20172841 (2018).

2. Kiltie, R. A. Distribution of palm fruits on a rain forest floor: why white-lipped peccaries forage near objects. Biotropica 14, 141–145 (1981).

3. Bodmer, R. E. Responses of ungulates to seasonal inundations in the Amazon floodplain. J. Trop. Ecol. 6, 191–201 (1990).

4. Vickers, W. T. Hunting yields and game composition over ten years in an Amazon Indian territory. Neotropical Wildlife Use and Conservation 400, 53–81 (1991).

5. Fragoso, J. M. V. Desapariciones locales del baquiro labiado (*Tayassu pecari*) en la Amazonía: migración, sobre-cosecha o epidemia. Manejo de Fauna Silvestre en la Amazonía (Fang, T., Bodmer, R., Aquino, R. & Valqui, M. eds.). United Nations Development Program-Global Environmental Facility, Universidad de Florida, Instituto de Ecología. La Paz, Bolivia, 309–312 (1997).

6. Fragoso, J. M. V. A long-term study of white-lipped peccary (*Tayassu pecari*) population fluctuation in northern Amazonia. People in Nature, Wildlife Conservation in South and Central America (Silvius, K., Bodmer, R. E. & Fragoso, J. M. V. eds.). Columbia University Press, New York, 286–296 (2004).

7. Ripple, W. J. et al. Collapse of the world’s largest herbivores. Science Adv. 1, e1400103 (2015).

8. Sobral, M. et al. Mammal diversity influences the carbon cycle through trophic interactions in the Amazon. Nat. Ecol. Evol. 1, 1670–1676 (2017).

9. Fragoso J. M. V. et al. Line transect surveys under detect terrestrial mammals: implications for the sustainability of subsistence hunting. PLoS ONE 11, e0152659 (2016).

10. Donkin, R. A. The peccary: with observations on the introduction of pigs to the New World. Trans. Am. Philos. Soc. 75, 1–152 (1985).

11. Fausto, C. Feasting on people: eating animals and humans in Amazonia. Curr. Anthropol. 48, 497–530 (2007).

12. Virtanen, P. K. The Death of the Chief of Peccaries: The Apurinã and the Scarcity of Forest Resources in Brazilian Amazonia. Hunter-gatherers in a Changing World (Reyes-Garcia, V. & Pyhälä, A. eds.). Springer, Cham, 91–105 (2017).

13. Constantino, P. A. L. et al. Indigenous collaborative research for wildlife management in Amazonia: the case of Kaxinawá, Acre, Brazil. Biol. Conserv. 141, 2718–2729 (2008).

14. Iwamura, T., Lambin, E. F., Silvius, K. M., Luzar, J. B., & Fragoso, J. M. V. Agent-based modeling of hunting and subsistence agriculture on indigenous lands: understanding interactions between social and ecological systems. Environ. Modell. & Softw. 58, 109–127 (2014).

15. Emmons, L. H. Comparative feeding ecology of felids in a neotropical rainforest. Behav. Ecol. Sociobiol. 20, 271–283 (1987).

16. Young, T. P. Natural die-offs of large mammals: Implications for conservation. Conserv. Biol. 8, 410–418 (1994).

17. Smith, K. F., Sax, D. F. & Lafferty, K. D. Evidence for the role of infectious disease in species extinction and endangerment. Conserv. Biol. 20, 1349–1357 (2006).

18. Fragoso, J. M. V. Home range and movement patterns of white-lipped peccary (*Tayassu pecari*) herds in the northern Brazilian Amazon. Biotropica 30, 458–469 (1998a).

19. Richard-Hansen, C. et al. Movements of White-Lipped Peccary in French Guiana. Movement Ecology of Neotropical Forest Mammals: Focus on Social Animals (Reyna-Hurtado, R. & Chapman, C. A. eds.). Springer, Cham, 57–75 (2019).

20. Fragoso, J. M. V. White-lipped peccaries and palms on the Ilha de Maracá. Maracá: The Biodiversity and Environment of an Amazonian Rainforest. J. Wiley & Sons, New York, 15–64, (1998b).

21. Richard-Hansen, C., Surugue, N., Khazraie, K., Le Noc, M. & Grenand, P. Long-term fluctuations of white-lipped peccary populations in French Guiana. Mammalia 78, 291–301, (2014).

22. Brocardo, C. R., da Silva, M. X., Delgado, L. E. & Galetti, M. White-lipped peccaries are recorded at Iguaçu National Park after 20 years. Mammalia 81, 519–522 (2017).

23. Antunes, A. P. et al. A conspiracy of silence: Subsistence hunting rights in the Brazilian Amazon. Land Use Policy 84, 1–11 (2019).

24. Ministerio del Ambiente MINAM. Dictamen de Extractión No Perjudicial (DENP) de los cueros de pecaríes (Pecari tajacu y Tayassu pecari) en el Perú. Ministerio del Ambiente, Dirección General de Diversidad Biológica. Lima, Perú. (2020).

25. Constantino, P. A. L. Deforestation and hunting effects on wildlife across Amazonian indigenous lands. Ecol. Soc. 21, 1–10 (2016).

26. Fragoso, J. M. V. Perception of scale and resource partitioning by peccaries: behavioral causes and ecological implications. J. Mammal. 80, 993–1003 (1999).

27. Forchhammer, M. C., Stenseth, N. C., Post, E. & Langvatn, R. Population dynamics of Norwegian red deer: density-dependence and climatic variation. Proc. R. Soc. Lond. B 265, 341–350 (1998).

28. Owen-Smith, N. Modeling the population dynamics of a subtropical ungulate in a variable environment: rain, cold and predators. Nat. Resour. Model. 13, 57–87 (2000).

29. Imperio, S., Focardi, S., Santini, G. & Provenzale, A. Population dynamics in a guild of four Mediterranean ungulates: density-dependence, environmental effects and inter-specific interactions. Oikos 121, 1613–1626 (2012).

30. Altrichter, M., et al. Range-wide declines of a key Neotropical ecosystem architect, the Near Threatened white-lipped peccary *Tayassu pecari*. Oryx 46, 87–98 (2012).

31. Krebs, C. J. Population cycles revisited. J. Mammal. 77, 8–24 (1996).

32. Ginzburg, L., & Colyvan, M. Ecological Orbits: How Planets Move and Populations Grow. Oxford University Press, New York (2004).

33. Eisenberg, J. F. & Redford, K. H. The contemporary mammalian fauna. Mammals of the Neotropics-the central neotropics, 3. University of Chicago Press, Chicago (1999).

34. Weaver, S. C. & Reisen, W. K. Present and future arboviral threats. Antiviral Research 85, 328–345 (2010).

35. Kissler, S., Tedijanto, C., Lipsitch, M. & Grad, Y. H. Social distancing strategies for curbing the COVID-19 epidemic. medRxiv https://doi.org/10.1101/2020.03.22.20041079 (2020).

36. McCallum, H., & Dobson, A. Disease and connectivity. Connectivity Conservation (Crooks, K. & Sanjayan, M. eds.). Cambridge University Press, Cambridge, 479–501. doi:10.1017/CBO9780511754821.022, (2006).

37. Juárez E. A. Y., Mace, G. M., Cowlishaw, G. & Pettorelli, N. Natural population die-offs: causes and consequences for terrestrial mammals. Trends Ecol. Evol. 27, 272–277 (2012).

## Online Methods References

38. Ospina, R. & Ferrari, S. L. Inflated beta distributions. Stat. Papers 51, 111 https://doi.org/10.1007/s00362-008-0125-4 (2010).

39. Stasinopoulos, M. D., Rigby, R. A., Heller, G. Z., Voudouris, V. & De Bastiani, F. Flexible Regression and Smoothing: using GAMLSS in R. CRC Press, Boca Raton, Florida (2017).

40. Hastie, T. J. & Tibshirani, R. J. Generalized Additive Models (Vol. 43). CRC press, Boca Raton, Florida (1990).

41. Fewster, R. M., Buckland, S. T., Siriwardena, G. M., Baillie, S. R. & Wilson, J. D. Analysis of population trends for farmland birds using generalized additive models. Ecology 81, 1970–1984 (2000).

42. Rigby, R. A. & Stasinopoulos, D. M. Generalized additive models for location, scale and shape J R Stat Soc Ser C Appl Stat 54, 507–554 (2007).

43. Dalgaard, P. (Producer) R Development Core Team. R: A language and environment for statistical computing. Computer programme http://www.R-project.org/ (2010).

