## Supplemental Materials for "Mysterious disappearances of a large mammal in Neotropical forests"

**Supplementary Table 1 References**


44. Azevedo, F. C. C. & Conforti, V. A. Decline of peccaries in a protected subtropical forest of Brazil: toward conservation issues. *Mammalia* **72**, 82-88 (2008).

45. Barbarán, F. Estado del hábitat y registros de la presencia del tigre (*Panthera onca*) en el área de influencia de la Reserva Provincial Acambuco (Provincia de Salta, Argentina). *Ecosistemas* **13**: 88-95 (2004).

46. Taber, A. et al. *El Destino de los Arquitectos de los Bosques Neotropicales: Evaluación de la Distribución y Estado de Conservación de los Pecaríes Labiados y los Tapires de Tierras Bajas*. IUCN/SSC Tapir Specialist Group and Peccary Specialist Group, IUCN, Wildlife Conservation Society & Wildlife Trust. 181 pp. (2008).

47. de Bustos, S. et al. Situación crítica del pecarí labiado *Tayassu pecari* en las Yungas de la Alta Cuenca del Río Bermejo, Argentina. *Libro de Resumen XXII Jornadas Argentinas de Mastozoología, Argentina*, Sociedad Argentina para el Estudio de los Mamíferos (2018).

48. Bardavid, S., de Bustos, S., Politi, N. & Rivera, L. Escasez de registros de pecarí labiado (*Tayassu pecari*) en un sector de alto valor de conservación de las Yungas australes de Argentina. *Mastozool. Neotrop.* **26**,167-173 (2019).

49. de Bustos, S., Varela, D., Lizárraga, L., Camino, M. & Quiroga, V.A.  Tayassu pecari. *Categorización 2019 de los mamíferos de Argentina según su riesgo de extinción. Lista Roja de los mamíferos de Argentina* (SAyDS–SAREM eds.). http://cma.sarem.org.ar (2019).

50. Perovic, P. Ecología de la comunidad de félidos en las Selvas Nubladas del Noroeste Argentino. Ph.D. dissertation. Fac. de Cs. Exactas, Físicas y Naturales, Universidad Nacional de Córdoba. 125 pp. (2002).

51. Stearman, A. M. Making a living in the tropical forest: Yuqui foragers in the Bolivian Amazon. *Hum. Ecol.* **19**, 245-260 (1991).

52. Stearman, A. M. & Redford, K. H. Game management and cultural survival: the Yuqui ethno development project in lowland Bolivia. *Oryx* **29,** 29-34 (1995).

53. Wallauer, J. P. & Albuquerque, E. P. Lista preliminar dos mamíferos de observados no Parque Estadual do Turvo, Tenente Portela, Rio Grande do Sul, Brasil. *Roessléria* **8**, 179-185 (1986).

54. Keuroghlian, A. et al. Avaliação do risco de extinção do queixada, *Tayassu pecari* (Link, 1795) no Brasil. *(Extinction risk assessment of white-lipped peccaries in Brazil). Biodiversidade Brasileira* **3**, 84-102 (2012).

55. Kasper, C. B., Mazim, F. D., Soares, J. B. G., Oliveira, T. G. & Fabian, M. E. Composição e abundância relativa dos mamíferos de médio e Grande Porte no Parque Estadual do Turvo, Rio Grande do Sul, Brasil. *Rev Bras Zool* **24**, 1087-1100 (2007).

56. Beisiegel, B., Nakano, E. & Jorge, M. L. S. P. Are white-lipped peccaries back in the Paranapiacaba Forest, São Paulo, Brazil? *Suiform Soundings* **12**, 29-33 (2014).

57. Beisiegel, B. M. Shelter availability and use by mammals and birds in an Atlantic forest area. *Biota Neotropica* **6**, 1-16 (2006).

58. Rocha, D. G. Padrão de Atividade e Fatores que Afetam a Amostragem de mamiferos de

Medio e Grande Porte na Amazonia. MSc. Dissertation. Instituto Nacional de Pesquisas da Amazônia Central (2015).

59. Fragoso, J. M. V. Large Mammals and the Community Dynamics of an Amazonian Forest. Ph.D. Dissertation, University of Florida, USA. (1994).

60. Borges, L. H. M., Calouro, A. M., & Sousa J. R. Large and medium-sized mammals from Chandless State Park, Acre, Brazil. *Mastozool. Neotrop.* **22**, 265-277 (2015).

61. Hallett, M. T. et al. Impact of low-intensity hunting on game species in and around the Kanuku Mountains Protected Area, Guyana. *Front. Ecol. Evol.* **7**, 412 (2019).

62. Hallett, M. T. Landscape-scale research as a tool for engaging communities in a shared learning process for conservation and management in the Rupunuini, Guyana, Ph.D. Dissertation, University of Florida. https://ufdcimages.uflib.ufl.edu/UF/E0/05/18/39/00001/HALLETT_M.pdf (2017).

63. Paemelaere, E. A. D., Fernandes, D., Leroy, I. & Angelbert, J. Large Mammals of the South Rupununi Region, Guyana. *Biodiversity Assessment Survey of the South Rupununi Savannah, Guyana*. (Alonso, L.E., Persaud, J. & Williams, A. eds.). BAT Survey Report No. 1. WWF-Guianas, Guyana Office. Georgetown, Guyana, 119-134 (2016).

64. Roopsind, A., Caughlin, T. T., Sambhu, H., Fragoso, J. M.V. & Putz, F. E. Logging and indigenous hunting impacts on persistence of large Neotropical animals. *Biotropica* **49**, 565-575 (2017).

65. Shaffer, C. A., Milstein, M. S., Yukuma, C., Marawanaru, E. & Suse, P. Sustainability and comanagement of subsistence hunting in an indigenous reserve in Guyana. *Biol. Conserv.* **31**, 1119-1131 (2017).

66. Kiltie, R. A. & Terborgh, J. Observations on the behavior of rain forest peccaries in Perú: Why do white-lipped peccaries form herds? *Z. Tierpsychol.* ***62***, 241-255 (1983).

67. Silman, M. R., Terborgh, J. & Kiltie, R. Population regulation of a dominant-rain forest tree by a major seed-predator. *Ecology* **84**, 431-438 (2003).

68. Mayor, P. et al. Effects of selective logging on large mammal populations in a remote indigenous territory in the northern Peruvian Amazon. *Ecol. Soc.* **20**, 36 (2015).
